## Supplementary figures and images for "A SARS-CoV-2 vaccine designed for manufacturability results in unexpected potency and non-waning humoral response"

### STable 1

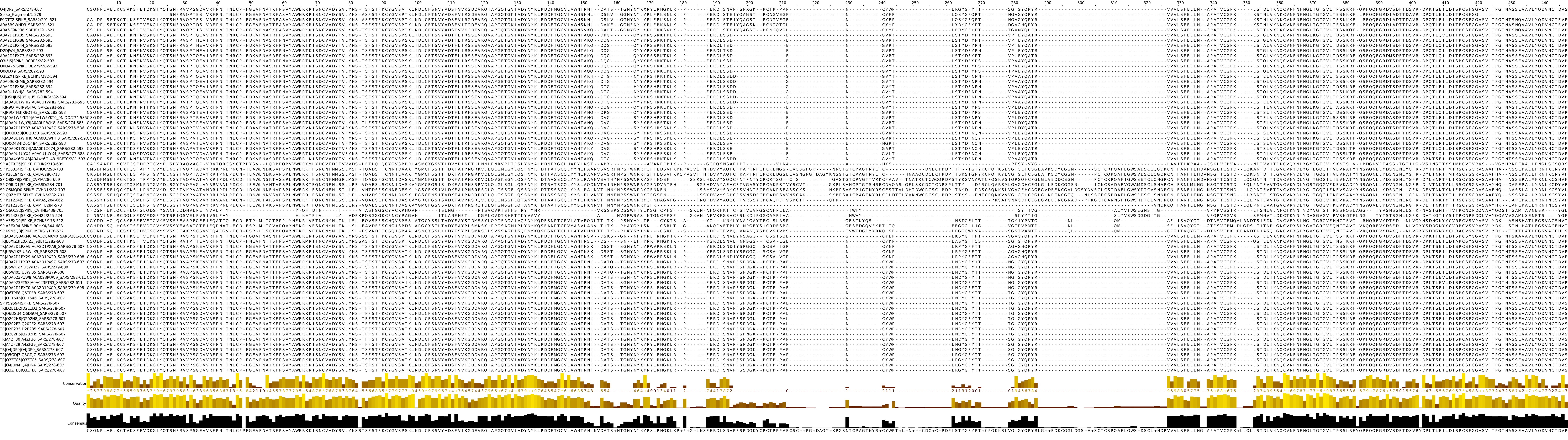
